# TOR acts as metabolic gatekeeper for auxin-dependent lateral root initiation in *Arabidopsis thaliana*

**DOI:** 10.1101/2022.03.29.486207

**Authors:** Michael Stitz, David Kuster, Maximilian Reinert, Mikhail Schepetilnikov, Béatrice Berthet, Denis Janocha, Anthony Artins, Marc Boix, Rossana Henriques, Anne Pfeiffer, Jan Lohmann, Emmanuel Gaquerel, Alexis Maizel

## Abstract

Plants post-embryonic organogenesis requires matching the available metabolic resources to the developmental programs. The root system is determined by the formation of lateral roots (LR), which in *Arabidopsis thaliana* entails the auxin-induced activation of founder cells located in the pericycle. While the allocation of sugars to roots influences root branching, how sugar availability is sensed for auxin-triggered formation of LRs remains unknown. Here, we combine metabolic profiling with cell-specific genetic interference to show that LR formation is an important sink for carbohydrate accompanied by a switch to glycolysis. We show that the target-of-rapamycin (TOR) kinase is locally activated in the pericycle and the founder cells and that both chemical and genetic inhibition of TOR kinase lead to a block of LR initiation. TOR marginally affects the auxin-induced transcriptional response of the pericycle but modulates the translation of ARF19, ARF7 and LBD16, three key targets of auxin signalling. These data place TOR as a gatekeeper for post-embryonic LR formation that integrates local auxin-dependent pathways with systemic metabolic signals, modulating the translation of auxin induced gene expression.

## Introduction

Plants assimilate atmospheric CO_2_ in their leaves and convert it into simple sugars by photosynthesis. Sucrose is the predominant sugar transported from the source tissues to heterotrophic sink tissues where it is hydrolysed to fructose and glucose that fuels growth and development. The root system is an obligate sink organ and mounting evidence suggests that allocation of sugars to roots drives primary root growth and lateral root development, two main determinants of the root system architecture. Shortly after germination, light triggers root growth via the transport of photosynthesis-derived sugar into the root tip (Kircher & Schopfer, 2012; Xiong *et al*, 2013; Yuan *et al*, 2014) and increased photosynthetic rates in above-ground tissues correlate with increased lateral root (LR) formation (Crookshanks *et al*, 1998). Crosstalk between carbon metabolism and phytohormone signalling, mainly auxin signalling, has been linked to the modulation of root system architecture (Sairanen *et al*, 2012; Lilley *et al*, 2012; Gupta *et al*, 2015). Regardless of the known role sugar plays in lateral root development, our knowledge about how sugars and glucose modulate LR formation at the molecular level remains unknown.

LR formation is an auxin-controlled process which in *Arabidopsis thaliana* (hereafter *Arabidopsis*), occurs through activation of founder cells that undergo a series of cell divisions to form a primordium that emerges from the primary root (Malamy & Benfey, 1997). Founder cells are located in the pericycle facing the xylem pole and the earliest marker of LR initiation is their radial swelling, repolarization and nuclei migration towards the common anticlinal wall (Schütz *et al*, 2021; Vilches Barro *et al*, 2019; von Wangenheim *et al*, 2016). Additional founder cells are recruited (Torres-Martínez *et al*, 2020) which further proliferate and form a LR dome-shaped primordium (LRP) (Lucas *et al*, 2013). Upregulation of auxin signalling (Dubrovsky *et al*, 2008) and of *GATA23* expression (De Rybel *et al*, 2010) are two molecular markers associated with LR founder cells and initiation. Auxin-dependent gene regulation plays a major role in all stages of LR development and occurs through TRANSPORT INHIBITOR RESISTANT 1/AUXIN SIGNALLING F-BOX (TIR1/AFB) induced degradation of the AUXIN/INDOLEACETIC ACID (Aux/IAA) repressors that frees the transcriptional activators AUXIN RESPONSE FACTORS (ARFs) inducing expression of downstream genes (Blázquez *et al*, 2020). During LR initiation, Aux/IAA 14 (*IAA14, SOLITARY ROOT*), *ARF7* and *ARF19* are necessary for cell cycle entry (Fukaki *et al*, 2002; Okushima *et al*, 2005; Wilmoth *et al*, 2005) and activation of LATERAL ORGAN BOUNDARY (LBD) 16, a transcription factor, is required for the asymmetric division of these cells (Okushima *et al*, 2007; Goh *et al*, 2012). The cell and mechanistic bases of lateral root initiation start to be elucidated (Santos Teixeira & Ten Tusscher, 2019). However, insight into how the plant’s metabolic status is integrated in the regulation of this developmental program is mostly unknown.

Energy availability perception is mediated in plants by two evolutionarily conserved and counteracting kinases (Shi *et al*, 2018; Crepin & Rolland, 2019). The SUCROSE NON FERMENTING1 RELATED PROTEIN KINASE1 (SnRK1) promotes catabolic metabolism, contrasting the TARGET OF RAPAMYCIN (TOR) kinase that promotes anabolic, energy-consuming processes. TOR forms a complex (TORC), consisting of TOR, and the TOR-interacting proteins RAPTOR (regulatory-associated protein of mTOR, RAPTOR1A and RAPTOR1B) and LST8 (small lethal with SEC13 protein 8, LST8-1 and LST8-2 (Menand et al., 2002; Anderson et al., 2005; Deprost et al., 2005; Moreau et al., 2012)). While *tor*-null mutants are embryonic lethal (Menand *et al*, 2002), the predominantly expressed regulatory proteins RAPTOR1B and LST8-1 show viable mutant phenotypes (Salem *et al*, 2017; Moreau *et al*, 2012). TORC is activated by nutrients (Dobrenel *et al*, 2016) such as glucose and branched-chain amino acids (Cao *et al*, 2019) as well as phytohormones such as auxin (Schepetilnikov *et al*, 2017) and it phosphorylates targets linked to cell cycle, translation, lipid synthesis, N assimilation, autophagy and ABA signalling (Shi *et al*, 2018; Van Leene *et al*, 2019). The usage of inducible TOR knockdown lines (Xiong *et al*, 2013) and specific chemical inhibitors like AZD8055 (Montané & Menand, 2013), led to the discovery of a mechanistic connection between TOR and its phosphorylation substrates to a multitude of developmental processes (Van Leene *et al*, 2019; Shi *et al*, 2018). In particular, TOR is essential for the activation of the embryo-derived root and shoot meristems during the photoautotrophic transition (Xiong *et al*, 2013; Pfeiffer *et al*, 2016; Li *et al*, 2017). While TOR plays a central role in coordinating the energy-status of plants with several developmental programs from embryo development to senescence (Shi *et al*, 2018), it remains unknown whether TOR plays a role in the post-embryonic establishment of new meristems like in the case of LR formation. Two reports suggest a possible link between energy availability sensing pathways and LR formation. Using an engineered rapamycin sensitive version of TOR in potato, it was shown that TOR is necessary for hypocotyl-borne (adventitious) root formation (Deng *et al*, 2017) and, recently, SnRK1 was shown to be required for LR formation induced by unexpected darkness (Muralidhara *et al*, 2021).

Here, we examine the role of TOR in LR formation and its interplay with auxin signalling. Using a combination of root metabolomics profiling, sugar metabolism manipulation, chemical/genetic and tissue specific inhibition of TOR-dependent signalling, and genome wide profiling of TOR effects on transcriptome and translatome, we present evidences that TOR activation in the pericycle and LR founder cells and subsequent LR initiation is tuned by high glycolysis rates depending on shoot-derived sugar. We further show that TOR marginally influences the auxin-induced transcriptional response of the pericycle but rather modulates the translation of several auxin response genes such as ARF19, ARF7 and LBD16. Altogether these data support a model placing TOR as a gatekeeper for post-embryonic LR formation that integrates systemic metabolic signals and local development by modulating the translation of auxin induced gene expression.

## Results & Discussion

### Plants with impaired lateral root formation hyperaccumulate starch in foliage

LR formation is an energy demanding process that depends on shoot derived carbon. Photoassimilates not consumed immediately by the plant’s foliar metabolism are stored as transitory starch granules and consumed in the dark period till onset of the light period (Graf *et al*, 2010). We first set out to assess how LR carbon demand impacts on foliar carbon metabolism by performing starch staining in seedlings impaired in different steps of the auxin signalling cascade specifically controlling LR formation (Fig. 1A). At the end of the dark period, wild type Col-0 seedlings accumulated limited amounts of starch indicated by the characteristic purple stain (Fig. 1B). Globally impairing auxin signalling in LR-less *iaa14*/*solitary root* (*slr*, Fig. 1C) and *arf7/arf19* (Fig. 1D) mutant plants however resulted in intense blue coloration throughout the leaves, indicative of higher levels of starch accumulation. ARF7 and ARF19 regulate the transcriptional activation of *LBD16* and plants expressing a dominant repressor version of LBD16 (*LBD16-SRDX*, Fig. 1E) do not form LR (Goh *et al*, 2012). Starch staining in LR-less *LBD16-SRDX* seedlings, led to comparably intense dark coloration as observed for *slr* foliage, indicating that blocking LR formation by interfering with the auxin signalling cascade or its direct targets leads to hyperaccumulation of photoassimilates in leaves. To ascertain that this increased accumulation of starch was caused by the lack of LRs and not a systemic effect of interfering with auxin signalling, we performed starch staining in *pGATA23::shy2-2-GR* and *pGATA23::slr1-GR* plants. These lines express dominant repressor versions of *SLR* and *SHY2/IAA3* specifically in the pericycle upon application of dexamethasone (DEX), resulting in an inducible, pericycle specific inhibition of auxin signalling and LR formation (Ramakrishna *et al*, 2019). Whereas control-treated lines showed faint purple coloration in leaves comparable to that in Col-0 plants (Fig. 1F, G), upon DEX treatment we observed intense starch accumulation in the leaves indicating that, similar to the global effect observed in the *slr, arf7/19* and *gLBD16-SRDX* mutants, blocking LR formation specifically in the root pericycle is sufficient to induce starch hyperaccumulation in foliage (Fig. 1H-J). Taken together, these observations point towards LR formation and its concomitant resource consumption to drive an increase in demand for shoot-derived carbon sources, in agreement with starch being a major integrator of plant growth regulation (Sulpice *et al*, 2009).

**Fig. 1).**
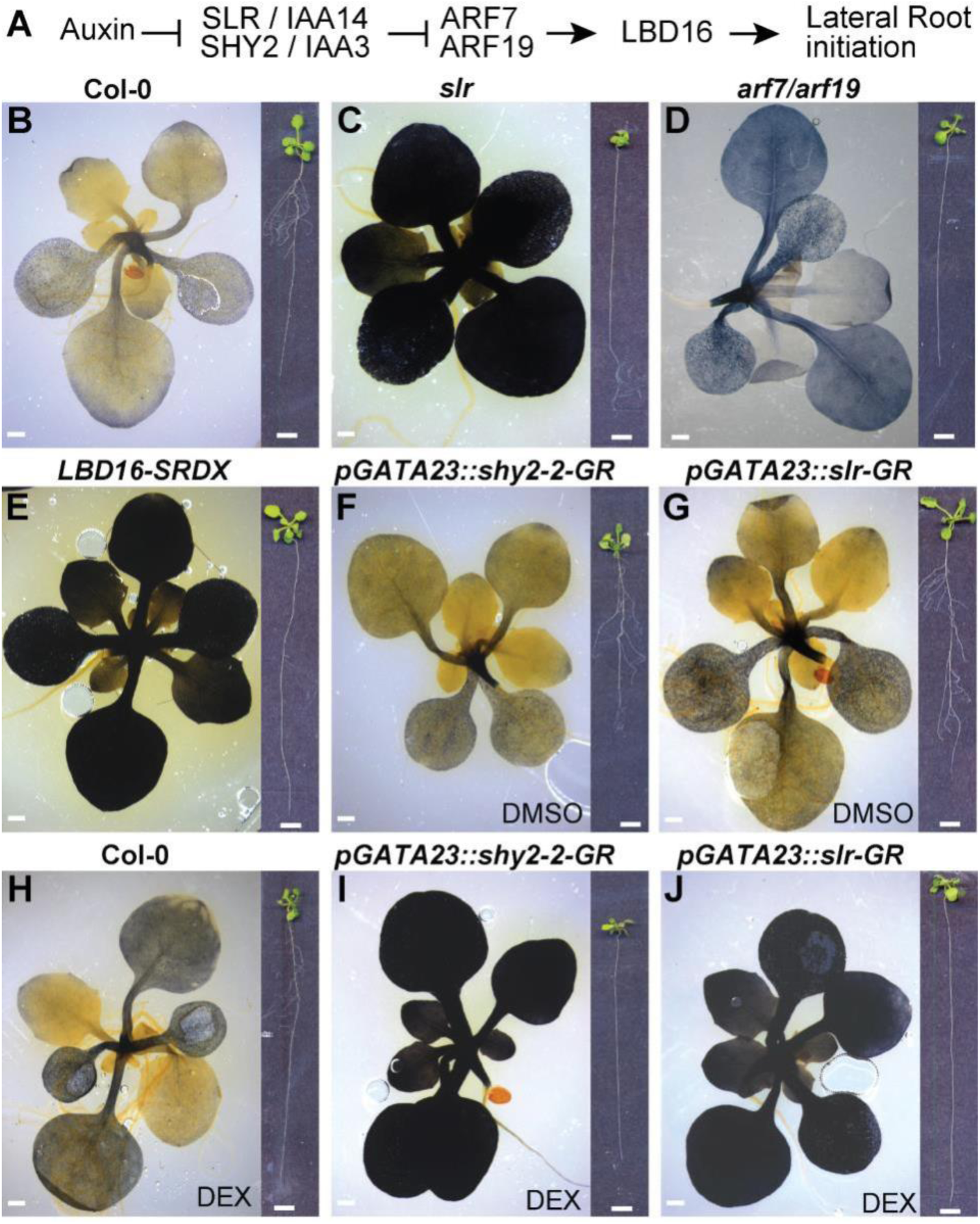
Lateral root deficiency leads to starch hyperaccumulation in leaves. **A**) Schematic representation of the auxin signalling module acting during lateral root initiation. **B-J**) Representative images of rosettes of 14-day-old seedlings stained with a Lugol’s Iodine solution for starch accumulation in Col-0 (**B**), *slr* (**C**), *arf7/arf19* (**D**) and *gLBD16-SRDX* (**E**) as well as in the inducible lateral root less lines *pGATA23::shy2-2-GR* and *pGATA23::slr1-GR* and Col-0 (**F-J**) grown on DMSO control medium (**F, G**) or on 30 µM Dexamethasone (DEX, **H-J**). Insets on the right show the root system. Scale bars: 0.5 cm.

### Auxin-triggered lateral root formation depends on shoot-derived carbohydrate catabolism in the root

To monitor changes in central carbon pathways resulting from the metabolism of shoot-derived carbohydrates and associated with the formation of LR, we conducted non-targeted metabolomics by gas chromatography coupled to mass spectrometry from shoot and root samples collected after synchronous induction of LR formation (Himanen *et al*, 2002). Briefly, after a pre-treatment with the auxin transport inhibitor NPA (N-1-naphthylphthalamic acid, 10 µM for 24h) that prevents LR initiation, 7-day-old seedlings were shifted to a medium containing auxin (Indole-3-acetic acid, IAA 10 µM) to synchronously activate the entire pericycle. Shoot and root samples were dissected and collected at six time points (0, 2h, 6h, 12h, 24h, 30h) after transferring seedlings from NPA to IAA to induce LR formation or maintaining them on NPA as control (Fig. 2A). This time series covers LR formation from initiation to stage V. By applying this procedure to wild type (Col-0) and *slr* mutant seedlings, we aimed at inferring a metabolic signature specifically associated with early stages of LR formation. To this end, raw metabolomics data were deconvoluted to extract compound-derived mass spectra used for annotation and statistical analysis. From a pool of more than 400 deconvoluted spectra, we conducted a hierarchical clustering analysis of the top 250 ones that exhibited non-constant intensity levels across the genotype x treatment x time matrix (Fig. 2B and File S1). This clustering analysis revealed that IAA induced several phases of reconfigurations of the root carbon metabolism in Col-0, which were largely altered in the *slr* mutant. Most specifically, cluster #1 comprised IAA-responsive compounds that were characterised in Col-0 roots by a slow build-up rate (reaching maximum values at 30h post IAA), the latter response was mostly impaired in the *slr* mutant (Fig. 2B). Cluster #2 and a sub-part cluster #4 were characterised by IAA-responsive compounds which reached much greater relative levels in the *slr* mutant 12h after transfer to IAA than in Col-0 roots (Fig. 2B).

**Fig. 2).**
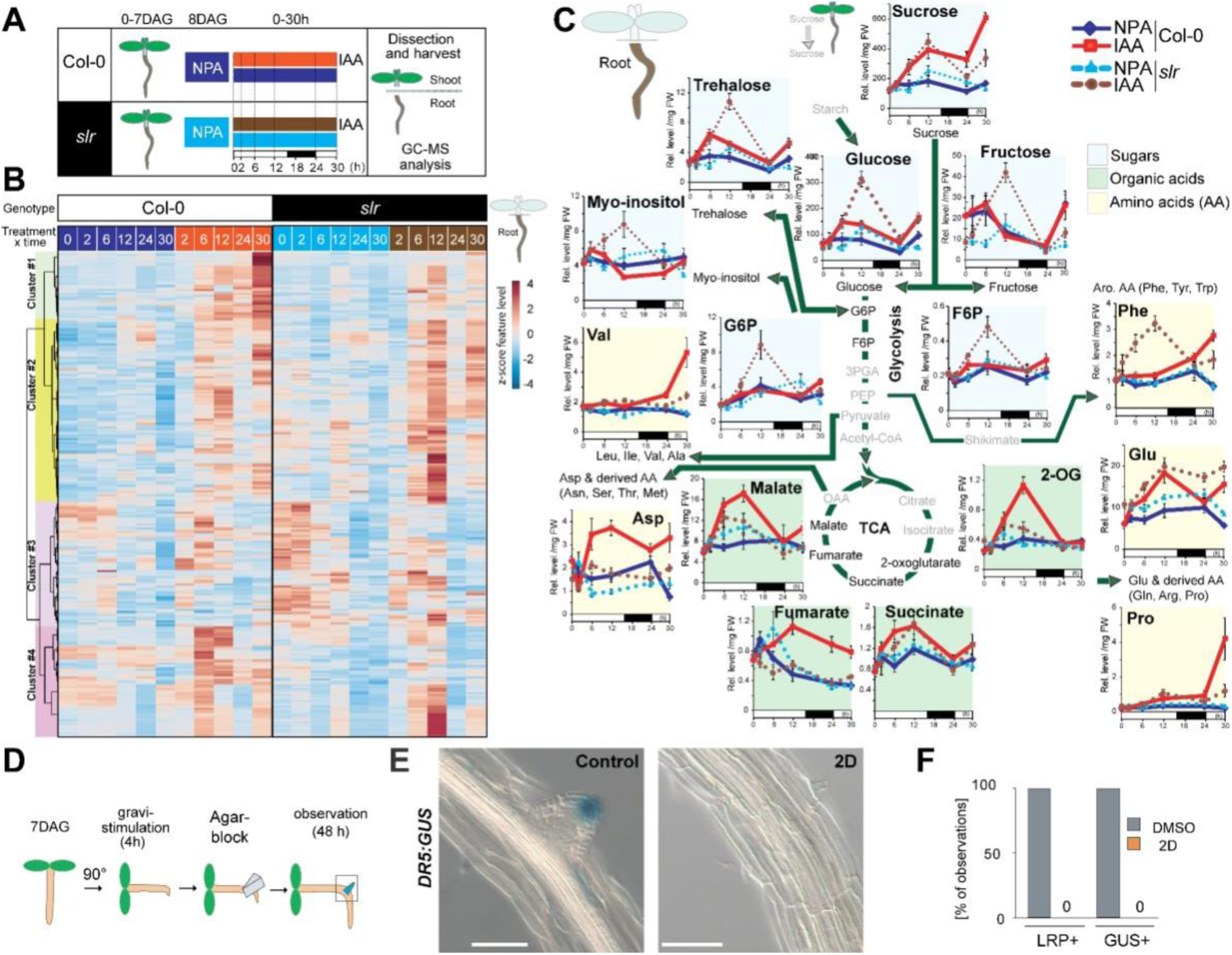
Increased flux within sugar glycolytic catabolism and connected pathways precedes and is essential for LR formation. **A**) Schematic of the experimental setup used for the GC-MS-based metabolomics profiling. **B**) Heatmap from a hierarchical clustering analysis (HCA) with Ward’s linkage showing z-score normalised relative levels of top 250 most intense compound-derived spectra (File S1) exhibiting non-constant intensity (One-way ANOVA & FDR-adjusted P<0.05) across experimental conditions in roots. Main HCA clusters are colour labelled. **C**) Mean relative levels (± SE, *n*=5, normalised to the ribitol internal standard and per mg fresh weight) for representative metabolites of sugar, glycolytic, tricarboxylic acid, amino acid metabolic pathways in root tissues of Col-0 (solid lines) and *slr* (dashed lines) at the indicated time after IAA application. White and black boxes below the x-axis indicate light and dark phases, respectively, during the sampling. Statistical differences for genotype x treatment (NPA-vs IAA-treated roots) are summarised in File S1. **D)** Schematic of the experimental setup for induction of LR formation upon local 2-deoxy-D-glucose (2D, 10mM) treatment. **E**) Representative differential interference contrast (DIC) images of root bends in DR5:GUS seedlings treated as indicated. 2D-treated bends did not develop LR primordia 48h after gravistimulation, scale bar: 50 µm. **F**) Fraction of root bends forming a LRP and showing DR5 GUS staining, after treatment with either control or 2D containing agar blocks, n=4.

We next mined clusters for metabolites associated with these IAA-/*slr*-dependent root metabolome responses. In line with the starch stainings data, levels of glucose and sucrose were slightly higher in *slr* shoot tissues compared to Col-0 (Fig. S1). Upon shift to IAA, levels of shoot sucrose quickly increased within two hours and remained high in Col-0 while it declined over the course of the day in control treated samples and remained mostly unchanged to presence of IAA in the *slr* mutant. This and the starch hyperaccumulation in shoots upon LR impairment indicate that LR formation relies on a sucrose transfer from the shoot. In seedling, photosynthetically derived sucrose has been described to act as an interorgan signal and as fuel to drive primary root growth (Kircher & Schopfer, 2012). For up to 24h after shift to IAA, sucrose levels built up similarly in root tissues of both Col-0 and *slr* and then became significantly higher in Col-0 than *slr* (Fig. 2C), in line with a weaker sink strength of *slr* resulting from its inability to form LRs. We looked at the levels of additional sugars and glycolytic intermediates which were found, from the clustering, to be deregulated in *slr* roots (Fig. 2B, C). Strikingly, root levels of glucose and fructose derived from sucrose cleavage, which did not build up in Col-0 upon IAA probably due to their catabolism by glycolysis, were strongly increased by the IAA treatment in the *slr* mutant (Fig. 2C). A similar IAA-dependent over-accumulation was detected for several additional sugars enriched within the IAA-regulated cluster visible at 12h in *slr* (Fig. 2B), such as the disaccharide trehalose, the polyol *myo*-inositol as well as glucose-6-P and fructose-6-P. Notably, glucose-6-P is produced by the hexokinase1 (HXK1) which when mutated was reported to reduce LR formation (Gupta *et al*, 2015) supporting that Glucose-6-P levels are instrumental for LR formation, a notion further supported in a recent study that showed that WOX7, a WUSCHEL-related transcription factor, acts downstream of HXK1 to regulate LR formation (Li *et al*, 2020). Trehalose 6-Phosphate (T6P) is an intermediate in trehalose formation and T6P has been proposed to serve as a signal to regulate sucrose metabolism and impinge on a range of developmental processes (Figueroa & Lunn, 2016). Whereas our analysis does not allow us to quantify the amount of T6P, the elevated levels of trehalose suggest that T6P levels may be also affected and potentially could influence LR formation.

The accumulation of glucose-6-P and fructose-6-P, two glycolysis intermediates, was in contrast with their normally low steady-state abundance (e.g. in Col-0 NPA/IAA conditions) characteristic of their rapid consumption by the glycolytic flux (Arrivault *et al*, 2009). Interestingly, soluble carbohydrates have been demonstrated to promote auxin biosynthesis (Sairanen *et al*, 2012). The accumulation of carbohydrates in roots when LR formation is compromised could thus explain the previously reported elevated levels of auxin observed in *slr* mutants (Vanneste *et al*, 2005). Downstream in the carbohydrate catabolic pathway, the prolonged increases in the levels of several intermediates of the tricarboxylic acid (TCA), observed in Col-0 but not in *slr*, indicate that auxin-induced LR formation increases the catabolic flux. Levels of several amino acids whose biosynthetic pathways connect to the TCA cycle further indicated that an up-regulation of energy-releasing and amino acid production pathways, previously reported at the transcriptomic level (Dembinsky *et al*, 2007) (Fig. S2) and here backed up by metabolite data, underpins early stages of LR formation and is impaired in *slr*. Together these data indicate that LR formation is associated with a switch to glycolysis. This observation echoes similar ones made in animals where acquisition of pluripotency has been linked to a switch to glycolysis supporting the concept of metabolic reprogramming of cell fate (Shyh-Chang & Ng, 2017).

To verify whether this activation of sugar usage through glycolysis is indeed required for LR formation and to rule out an effect of exogenous auxin, we induced LR formation by gravistimulation (Lavenus *et al*, 2015) in presence of 2-deoxy-d-glucose (2D) a non-metabolisable glucose analog blocking glycolysis. 2D was applied locally by an agar block positioned over the root bend (Fig. 2D). Whereas in all mock treated root bends a LR primordium, visualised with the *DR5:GUS* reporter, was visible, none could be observed when treated by 2D (Fig. 2D-F), indicating that carbohydrate metabolism is a prerequisite for LR formation.

Together these data show that glycolysis and glycolysis-dependent metabolic activations are required to form a LR.

### The TOR complex is activated upon lateral root induction

LR formation is an auxin induced process that is regulated by glucose (Gupta *et al*, 2015) and requires carbohydrate catabolism (our results). As the TOR complex (TORC) has been reported to be activated by glucose and auxin and to be required for root meristem activation (Xiong *et al*, 2013), we hypothesised that LR induction could lead to TORC activation. To test this, we monitored the phosphorylation of the canonical TORC substrate S6K1 in roots upon treatment with sucrose and IAA (Fig. 3A, B). Treatment with either sucrose or IAA led to an upregulation of S6K1 phosphorylation while co-treatment had a synergistic effect that was fully suppressed by treatment with the TORC inhibitor AZD8055 (Montané & Menand, 2013). Treating roots with IAA led to a glycolytic switch (Fig. 2B, C), we thus sought to check whether IAA effects on TORC activation are dependent on glycolysis. For this, we repeated the sucrose and IAA treatments in the presence of 2D. Inhibition of glycolysis led to a block of S6K1 phosphorylation induced by IAA indicating that in roots, TORC activation by auxin depends on carbohydrate metabolism. This result points to a difference in TOR behaviour in source (foliage) and sink tissues. Whereas in source tissues TOR activity is promoted by auxin (Schepetilnikov *et al*, 2017), our results indicate that in sink tissues such as the root, TOR activity is primarily promoted by auxin-induced promotion of sugar breakdown. This glycolysis-dependent promotion of TOR activity could be a specificity of heterotrophic tissues that allows a systemic integration of developmental progression with shoot photosynthetic capacity. Such coupling has been reported for the light-dependent regulation of alternative splicing in roots which is triggered by shoot-photosynthesized sugars and compromised when TOR levels are reduced or its activity reduced (Riegler *et al*, 2021).

**Fig. 3).**
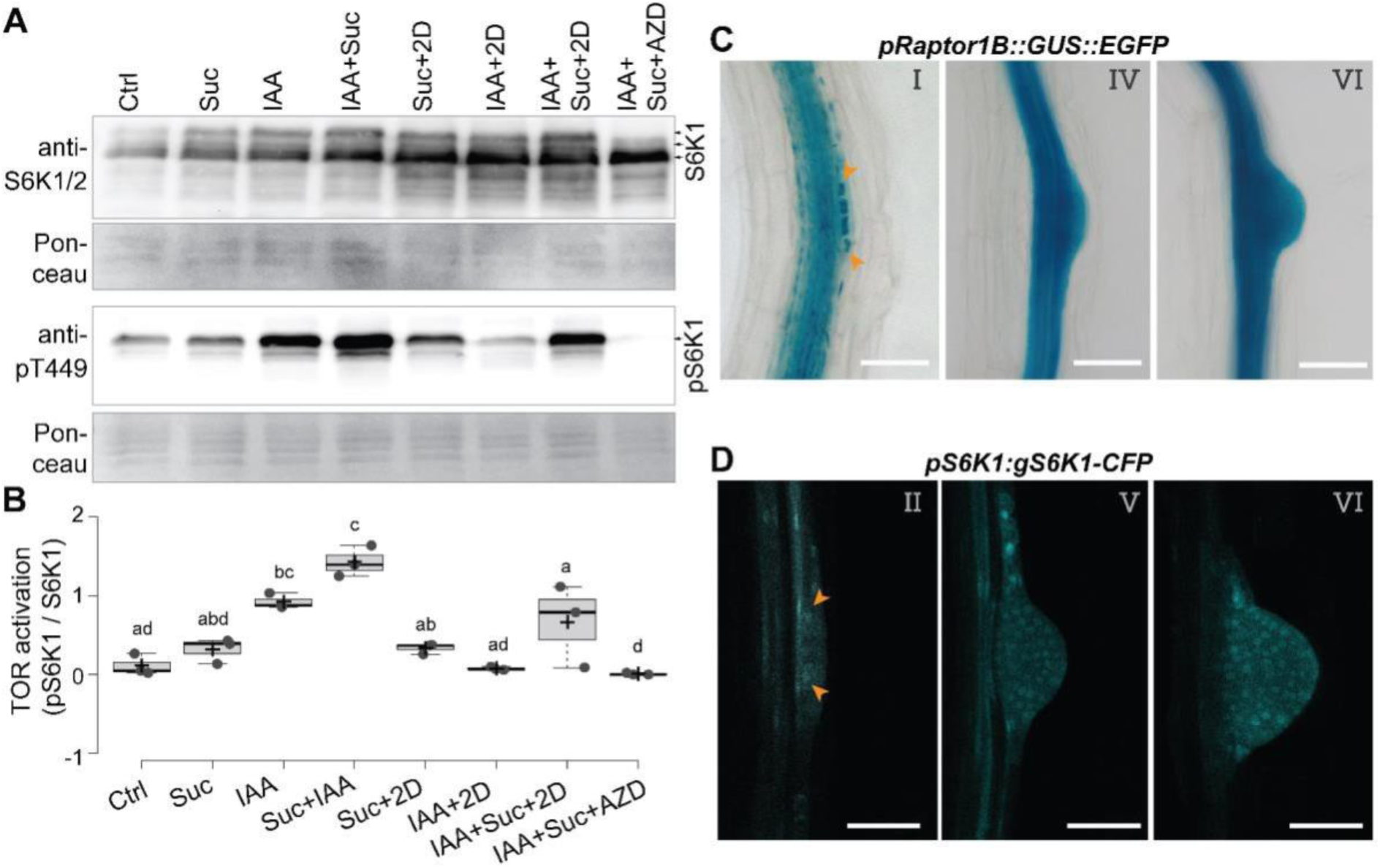
Auxin inducible S6K1 phosphorylation via TOR depends on activation of glycolysis in the primary root. **A)** Representative western blot of root tissues of *pUB10:S6K1-3xHA* treated by the indicated combination of auxin (IAA), sucrose (Suc), 2-deoxy-d-glucose (2D) and AZD8055 (AZD) and probed with anti-S6K1/2 or anti S6K1-T449P. Ponceau staining was used as loading control. **(B)** Quantification of the relative S6K activation. Box plots show three biological replicates and comparison between samples was performed by one-way ANOVA and post-hoc Tukey HSD Test (α = 0.05); different letters indicate significant differences. **C)** Representative DIC images showing RPT1B expression at different stages of LR development in 10 DAG *pRaptor1B::GUS::EGFP* seedlings. **D)** Representative confocal section showing S6K1 expression in different stages of LR development in 10 DAG *pS6K1:gS6K1-CFP* seedlings. Scale bars: 50 µm.

The previous results do not identify in which cells TOR is activated. To pinpoint in which tissues the TORC is present and active, we first used a reporter for the TORC subunit RAPTOR1B and detected its expression in the stele, LR founder cells of the pericycle and LR primordia (Fig. 3C). This expression pattern is similar to the one reported for the other TORC subunit LST8 (Moreau *et al*, 2012) and suggests that TORC is present in the forming LR. We also monitored in which cells S6K1 is expressed using a CFP-tagged genomic clone. This reporter specifically marked the actively dividing LR founder cells (Fig. 3D), confirming an earlier report (Zhang *et al*, 1994). Together these data suggest that the auxin-induced activation of glycolysis required for LR formation promotes the local activation of the TORC in the pericycle and the LR.

### The TOR complex is required for LR formation

To test whether TORC is necessary for the formation of LR, we looked at LR formation in plants with altered *TORC* levels or activity. We first used a *TOR* overexpression line (*TOR-oe*, (Deprost *et al*, 2007)) and observed longer primary roots and increased density of emerged LR indicating that elevated TOR levels can promote LR formation (Fig. 4A and S3). *TOR*-null mutants are embryo-arrested (Menand *et al*, 2002), we thus first quantified LR density in *raptor1b (rpt1b)*, a viable mutant affected in the scaffold protein that recruits substrates for TOR and leads to reduced TOR activity (Salem *et al*, 2017). LR density was reduced in *rpt1b*, indicating that the full TORC activity might be necessary for proper LR formation (Fig. 4B). To further confirm that TORC activity is required for LR formation, we treated wild type seedlings with AZD8055 (AZD) and conducted gravitropic stimulation (Fig. 4C). We found that AZD treatment led to a complete block of LR initiation (Fig. 4D). To test the contribution of *TOR* itself to LR formation, we designed a ß-estradiol (Est) inducible artificial miRNA against *TOR* that we first expressed from the *UBIQUITIN* promoter (*UB10pro>>amiR-TOR*). After 24h of Est treatment, *TOR* mRNA abundance was reduced to less than 25% of that of the DMSO control indicating efficient knock-down of *TOR* mRNA (Fig. S4). When LR formation was induced by gravistimulation in these conditions, we could not observe any division of the pericycle in the bend while stage III LR primordia were observed in DMSO treated plants (Fig. 4E). As exogenous application of sugar or auxin can promote LR formation and increased TOR activity, we checked whether the block in LR formation induced by knocking down *TOR* could be reversed by treatments with auxin and/or sucrose. Neither sole nor combined applications of sucrose or auxin could reverse the inhibition of LR formation induced by the *TOR* knockdown (Fig. S5). Together, reducing abundance or activity of TOR blocks LR formation at an early stage, indicating that TOR is essential. To determine whether *TOR* was required ubiquitously, or particularly in the LR founder cells, we specifically knocked down *TOR* in the xylem pole pericycle (XPP) cells from which LR founder cells derive (Parizot *et al*, 2008). For this, we drove the expression of the amiR-TOR from the XPP-specific promoter (Andersen *et al*, 2018; Vilches Barro *et al*, 2019) using a dexamethasone (Dex) inducible expression system (*XPPpro>>amiR-TOR*). We confirmed the tissue specificity of the *TOR*-knockdown in the *XPPpro>>amiR-TOR* line by starch stainings that reveal intense starch accumulation around the shoot vasculature in *XPPpro>>amiR-TOR*, a typical hallmark of impaired TORC function (Caldana *et al*, 2013) (Fig. S6). This excess in starch accumulation was also observed throughout the foliage of *UB10pro>>amiR-TOR* (Fig. S6). In the root, induction of *amiR-TOR* in the XPP cells led to a severe reduction in the number and density of LR formed compared to mock-induced plants (Fig. 4F,G) indicating that *TOR* is locally required in the pericycle to licence the auxin-induced formation of LR.

**Fig. 4).**
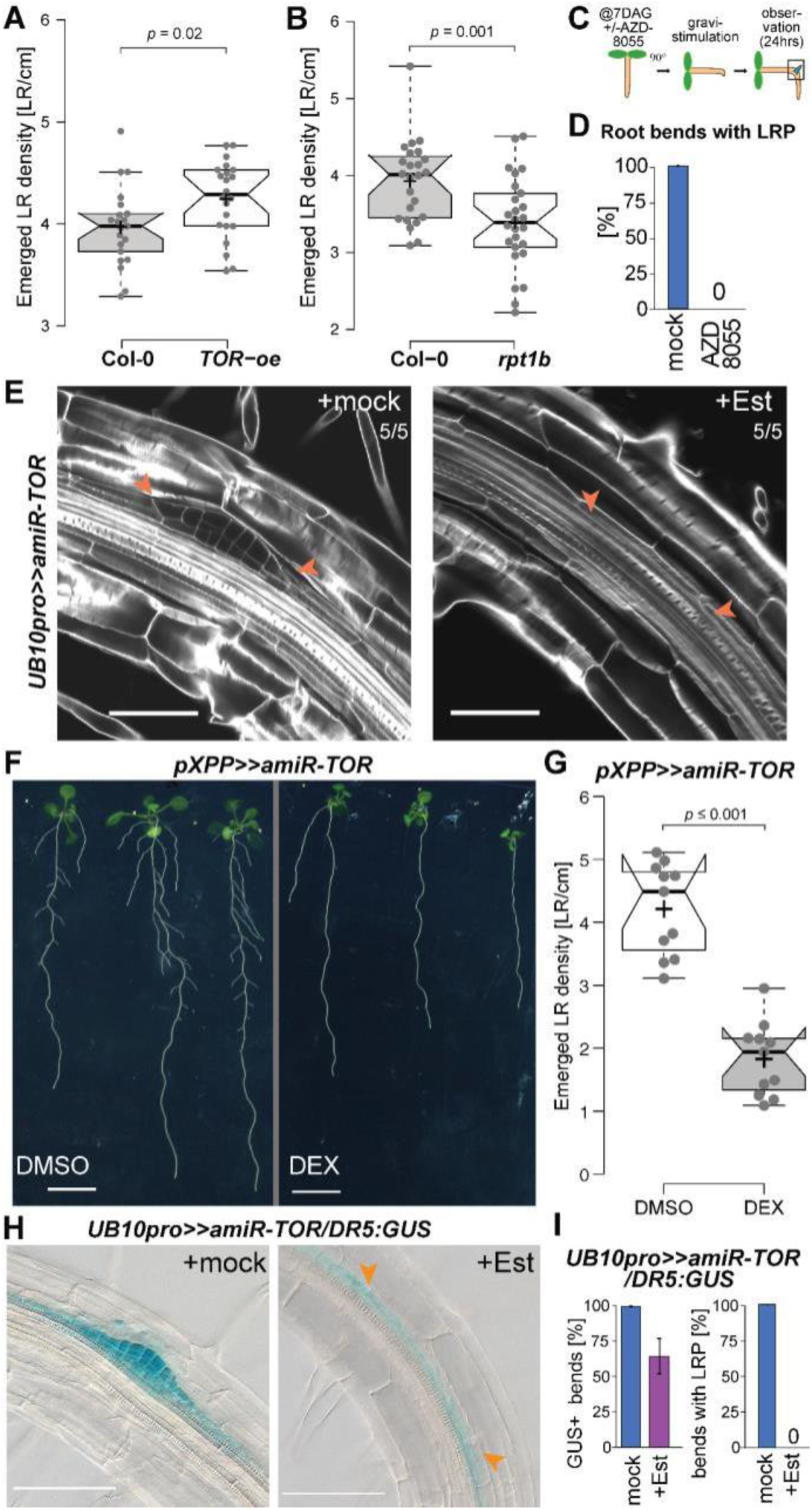
TORC is required in the pericycle for lateral root formation. Density of emerged LR in *TOR-oe* seedlings is increased, **(A)** and reduced in *rpt1b* mutant (**B**) when compared to Col-0 at 14 DAG. **C)** Schematic of the experimental setup used for scoring LR formation by gravistimulation upon inhibition of TOR by AZD8055. **D)** Proportion of bends developing lateral root primordia after transfer to AZD8055 containing media and gravistimulation for 24h (*n*=10). **E)** Representative confocal images of calcofluor counterstained bends of 7DAG *UB10pro>>amiR-TOR* seedlings following a 24 hr pre-treatment with mock (DMSO) or ß-Estradiol and subsequent 24h gravistimulation. Numbers indicate the penetrance of the phenotype. Scale bar: 50 µm. **F)** Phenotype of *pXPP>>amiR-TOR* seedlings grown on DMSO or Dexamethasone (DEX) at 14 DAG. Scale bar: 5mm. **G)** Density of emerged LR in *pXPP>>amiR-TOR* upon control or DEX treatment. **H)** Representative DIC images of bends in 7 DAG *UB10pro>>amiR-TOR/DR5*:*GUS* seedlings stained for GUS activity after a 24 hr pre-treatment with mock (DMSO) or ß-Estradiol (Est) and subsequent 24h gravistimulation, scale bar: 100 µm. **I)** Fraction of bends developing lateral root primordia and stained for GUS activity in primary root vasculature of *UB10pro>>amiR-TOR/DR5*:*GUS, n*=7.

Auxin accumulation in the XPP acts as a morphogenetic trigger for LR formation and is one of the earliest markers of LR initiation (Dubrovsky *et al*, 2008). To determine whether auxin accumulation was compromised in the *TOR* knockdown, we crossed the *pDR5::GUS* to the *UB10pro>>amiR-TOR* line. After gravistimulation for 24h, we observed *pDR5::GUS* accumulation in the apical region of the stage III LR-primordia in control conditions while upon *TOR* knockdown only a faint GUS-coloration was detected in the pericycle (Fig. 4H, I). Collectively, these data suggest that while *TOR* is required for LR initiation it does not compromise the formation of an auxin signalling maxima in the pericycle.

### TOR inhibition moderately affects the transcriptional auxin response associated with LR formation

While reduction of *TOR* abundance in the XPP or inhibition of its activity blocks LR formation, it does not block auxin signalling in these cells, suggesting that it is required either downstream or parallel to the auxin-induced LR formation developmental program. To get a genome wide picture of the effects of *TOR* knockdown during the early phase of LR formation, we compared by RNA-seq the transcriptomes of roots 6h after the synchronous induction of LR formation by auxin treatment (Himanen *et al*, 2002) in the inducible *UB10pro>>amiR-TOR* background (Fig. 5A). Transcriptome analysis identified 1141 auxin responsive genes in control conditions (Fig. S7). Upon *TOR* knock-down the expression of these genes was barely changed (Fig. 5B). We verified that inhibition of TOR activity by treatment with AZD8055 led to similar effects (Fig. S8). Collectively, although no morphological sign of LR initiation in the pericycle could be observed upon *TOR* knockdown or inhibition, the auxin-induced transcriptional response was globally unchanged. The SLR/IAA14 protein is a central regulator of LR initiation that controls the auxin-dependent expression of genes in the pericycle essential for LR initiation (Fukaki *et al*, 2002; Vanneste *et al*, 2005; Ramakrishna *et al*, 2019). We thus examined in detail how this set of IAA-induced genes behaved upon *TOR* knockdown. For this we took advantage of an existing dataset that profiles the response of the pericycle in similar conditions upon inhibition of *slr*-dependent auxin signalling in the pericycle (Ramakrishna *et al*, 2019) and identified 475 SLR-dependent genes responsive to auxin (Fig. S7). These genes behaved the same in *TOR* knockdown and in controls indicating that although LR formation is inhibited, the transcriptional SLR-dependent response to auxin in the pericycle is globally not affected when *TOR* levels are reduced. Together these data suggest that upon *TOR* reduction XPP cells still perceive and respond transcriptionally to auxin but appear unable to transform this response into a LR initiation event.

**Fig. 5).**
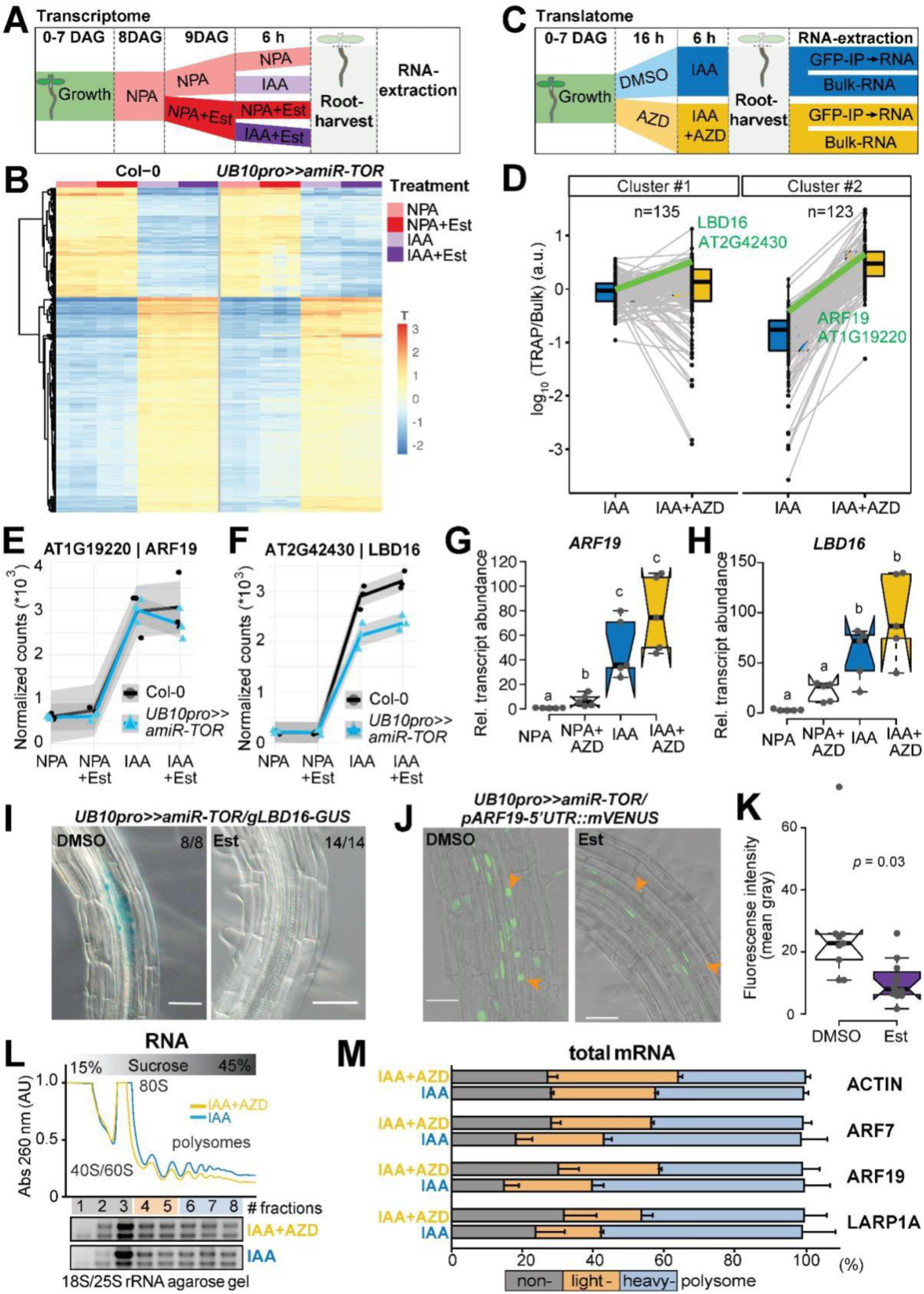
Effect of TOR on the auxin-induced lateral root transcriptome and translatome. **A)** Schematic of the experimental setup used to profile the impact of *TOR* knockdown on the transcriptome during LR formation. **B**) Heatmap from a k-means clustering analysis for 1141 IAA dependent transcripts (log fold change >1 & FDR <0.05). **C)** Schematic of the experimental setup used to profile the impact of *TOR* inhibition on the translatome during LR formation. **D)** Translational response (reads associated to ribosomes (TRAP) divided by reads in Bulk RNA) of 258 auxin induced genes. K-means revealed two clusters, with mild (#1) to strong (#2) shift in translation response upon TOR inhibition. The profiles of ARF19 and LBD16 are highlighted in green. **E, F)** Abundance of *ARF19* (E) and *LBD16* (F) transcripts in RNAseq samples. mRNA accumulation in response to auxin is comparable for both whether TOR is knocked down or not. **G, H**) Relative expression levels (normalised to *ACTIN*) of *ARF19* (G) and *LBD16* (H) measured by RT-qPCR upon TOR activity inhibition with AZD8055. Comparison between samples was performed by one-way ANOVA. Different letters indicate significant differences based on a post-hoc Tukey HSD Test (*n*=5, α = 0.05). **I)** Distribution of GUS-staining in *UB10pro>>amiR-TOR*/*gLBD16-GUS* seedlings 24 hrs after bending is absent if previously treated for 24 with Est (*n* = 8 -14). **J)** Representative confocal images of bends of 7 DAG *UB10pro>>amiR-TOR/pARF19-5’UTR::mVENUS* seedlings following a 24h pre-treatment with mock (DMSO) or ß-Estradiol and subsequent 24h gravistimulation. Scale bar: 50 µm, *n*=9. **K)** Signal (mean grey values) in the nuclei of the pericycle cells of *UB10pro>>amiR-TOR/ pARF19-5’UTR::mVENUS*. Significant differences between DMSO and Est-treated roots based on paired t-test, *n*=9. **L, M)** Total lysates prepared from lateral roots treated or not with IAA and AZD were fractionated through sucrose gradients, and the relative redistribution (percentage of total) of *ACTIN, ARF7, ARF19*, and *LARP1* mRNAs in each 8 fractions were studied by RT-qPCR analysis. (L) Polysome profiles. 40S, small ribosomal subunit; 60S, large ribosomal subunit; 80S, mono-ribosome; polysomes, polyribosomes. AU is arbitrary units of RNA absorbance at A260 nanometers. (M) RT-qPCR analysis of mRNA redistribution through sucrose gradient (8 fractions collected). Translation efficiency was computed as percentage of mRNA in non-polysome fractions (40/60/80S; fractions 1-3) against both light (fractions 4-5) and heavy polysomes (fractions 6-8). Plot is representative of two independently performed experiments with similar results. Data are mean +/- SEM.

### TOR affects translation of auxin-responsive transcription factors

The contrast between the mild effect of *TOR* knockdown on the root transcriptome and the strong block of LR formation, prompted us to investigate the effects of TOR inhibition on the translatome. To this end, we performed targeted purification of polysomal mRNA (TRAP-Seq, (Vragović *et al*, 2015)) using a transgenic line ubiquitously expressing a GFP-tagged RPL18 (Mustroph *et al*, 2009) 6h after the synchronous induction of LR formation by auxin treatment upon inhibition of TOR activity (AZD8055 treatment, AZD). To correct for abundance of mRNA, bulk RNA-Seq was performed on the same samples and used to normalise the reads purified with the ribosomes (Fig. 5C). This TRAP/Bulk ratio measures the fraction of mRNA associated with ribosomes, be it polysomes or monosomes and provides an indication of the degree of translation of a particular mRNA. Analysis of the bulk RNA-seq data identified 271 transcripts which are upregulated upon IAA treatment. Although this number is reduced compared to the transcriptome analysis due to absence of NPA pre-treatment, 80% of these genes were also differentially expressed in the *UB10pro>>amiR-TOR* transcriptome upon IAA treatment (Fig. S9). Clustering of these transcripts according to their TRAP/Bulk ratio between IAA and IAA+AZD conditions revealed two clusters. Cluster #1 consists of genes with moderate change in TRAP/Bulk ratio comparing IAA to IAA+AZD whereas the change was more important for cluster #2 (Fig. 5D). Examining these two clusters, we selected two candidates for further characterisation, LBD16 (cluster #1) and ARF19 (cluster #2), both involved in LR initiation (Okushima *et al*, 2007, 2005).

In the *UB10pro>>amiR-TOR* transcriptome, these two genes were induced by IAA and this induction was not affected by knocking down *TOR* (Fig. 5E, I). This result was independently confirmed by RT-qPCR upon inhibition of TOR activity by AZD8055 (Fig. 5F, J), indicating that transcription of these genes is not affected by TOR abundance or activity. In the translatome data, both genes had higher TRAP/Bulk ratio upon inhibition of TOR activity which suggest that the translation of these genes is different when TOR is inhibited. To verify this, we crossed the *UB10pro>>amiR-TOR* line to translational reporters for ARF19 and LBD16 and monitored the effect of *TOR* knock down on the expression of the reporters. For ARF19, the expression of the mVenus reporter is controlled by the ARF19 promoter, the 5’UTR and the 1st intron (*pARF19-5’UTR::mVenus*, (Truskina *et al*, 2021)). For LBD16, we used a GUS tagged genomic clone (Sheng *et al*, 2017). In both cases, whereas in control conditions expression of the reporters could be detected in the cells of the LR primordium, upon *TOR* knock down their expression was severely reduced while still present in the neighbouring cells (Fig. 5. G, H, K). This indicates that during LR initiation, TOR controls the expression of these two genes at the translational level. To further confirm that TOR can regulate the expression of genes at the level of translation, we looked at the association of the endogenous ARF19 transcript with ribosomes in wild type plants treated or not with AZD8055 during auxin-induced LR formation. Comparing the polysome profile upon TOR inhibition to the control revealed a shift from heavy to light fractions indicative of a reduced ribosomes processivity (Fig. 5L, M). This shift was very comparable to the one observed for *LARP1*, a transcript whose translation has been shown to be TOR dependent (Scarpin *et al*, 2020). Note that the distribution among ribosomal fractions was similar under both conditions for the housekeeping gene *ACTIN* (Fig. 5M). As ARF19 and ARF7 are jointly essential for LR initiation (Okushima *et al*, 2007), we also profiled the association of *ARF7* mRNA, whose expression is not auxin induced, with ribosomes. Like *ARF19, ARF7* mRNA shifted from heavy to light fractions upon TOR inhibition indicating that its translation is controlled by TOR (Fig. 5M). Together these data show that translation of both key transcription factors mediating auxin signalling during LR initiation is modulated by TOR. Intriguingly, the 5’UTR of both *ARF7* and *ARF19* mRNA contain several upstream open reading frames (uORF) that require a TOR-dependent translation re-initiation step to allow expression of the main ORF (Schepetilnikov *et al*, 2013) providing a likely mechanism by which TOR could regulate the expression of these genes. Collectively, our data support a model in which TOR acts as a metabolic gatekeeper for LR formation by locally integrating the availability of shoot derived photoassimilate with the auxin-mediated LR developmental program through control of the translation of key transcription factors. Such a model would ensure integration and coordination of the developmental and metabolic cues required for the formation of a new organ. Given the vast array of TOR outputs, it is likely that TOR may exert its gatekeeper role through additional mechanisms such as promotion of cell cycle progression via E2F as previously established (Xiong *et al*, 2013).

## Materials and Methods

### Plant material and growth conditions

Plants of *Arabidopsis thaliana* ecotype Colombia (Col-0) were grown under fluorescent illumination (50 µE m^-2^ s^-1^) in long day conditions (16 h light / 8 h dark) at 22 °C. Seeds were surface sterilised (ethanol 70% and SDS 0.1%) and placed on ½ Murashige and Skoog (MS) medium adjusted to pH 5.7 containing 1% agar (Duchefa). Following stratification (4°C in the dark, > 24 h). We used the previously described lines: *DR5::GUS* (Benková *et al*, 2003), *arf7/arf19* (Okushima *et al*, 2007), *LBD16-SRDX (Goh et al, 2012), slr* (Fukaki *et al*, 2002), *pGATA23::shy2-2-GR* and *pGATA23::slr1-GR* (Ramakrishna *et al*, 2019), *TOR-oe* (G548, (Deprost *et al*, 2007)), *raptor1b-1* (SALK_101990, (Salem *et al*, 2017), *35Spro::GFP-RPL18* (Mustroph *et al*, 2009), *pARF19-5’UTR::mVenus* (Truskina *et al*, 2021) and *gLBD16-GUS* (Sheng *et al*, 2017).

### Construction of vectors and plant transformation

Unless specified otherwise, the plasmids were generated using the GreenGate modular cloning system (Lampropoulos *et al*, 2013). For *pUB10:S6K1-3xHA*, the following modules were combined in pGGZ003: *UBQ10 promoter* (A), *B-dummy* (B), *S6K1* (AT3G08730, obtained by PCR on Col-0 gDNA (1398 bp) (C), *3xHA* (D), *35S terminator* (E) and *p35S:D-alaR:t35S* (F). The ß-Estradiol inducible amiR-TOR line (*UB10pro>>amiR-TOR)* was designed based on (Siligato *et al*, 2016) and two intermediate vectors (pAP039 and pAP043) were combined in pGGZ003. For pAP039 the following modules were combined in pGGM000: *pGGA044 Olex TATA* (A), *B-dummy* (B), TOR amiRNA (generated in this study, C), *D-dummy* (D), *RBCS* terminator (E) 250 bp HA adapter (G). For pAP043, the following modules were combined in pGGN000: *UBQ10 promoter* (A) *B-dummy* (B), CDS of chimeric *TF XVE* amplified from pLB12 (Brand *et al*, 2006) in two PCRs to domesticate an endogenous Eco31I site. (C), *D-dummy* (D), *UBQ10 terminator* (E) 250 bp HA adapter (G). For *pXPP::LhG4:GR/6xOP::amiR-TOR*, the intermediate module pAP097 was built consisting of *HA-adaptor, 6xOp* (A), *B-dummy* (B), *TORamiRNA* (C), *D-dummy* (D), *UB10* terminator (E) and HygrR (F) in pGGN000, and combined with pSW303 (in pGGM000) consisting of *pXPP* (A), *B-dummy* (B), *LhG4:GR* (C), *D-dummy* (D), *RBCS* terminator (E) FH-adaptor (F). Both modules were combined in pGGZ003 to generate the final vector. For the *pRAPTOR1B::GUS:eGFP* transcriptional fusion, 1360 bp upstream of the of *RAPTOR1B* (*AT3G08850*) were amplified by PCR, cloned into the pDONR221TM P1P2 by BP reaction (BP clonase, Thermofisher), and sub-cloned into the destination vector pHGWFS7.0 by LR reaction (LR clonase, Thermofisher). The *pS6K1:gS6K1-CFP* is a S6K1 genomic line with a C-terminal CFP clone. It was generated by PCR amplification (DNA KOD Hot-start DNA Polymerase, Novagen) and cloned into the pENTRD-TOPO Gateway vector using the manufacturer’s protocol and confirmed by sequencing. This clone was then used as templates to generate AscI-S6K1p::S6K1g(No STOP)-PacI fragments that were then ligated into the promoterless pBa002a vector to generate pBa002a/S6K1p::S6K1g-CFP. The clone was confirmed by sequencing. The primers used for cloning and sequencing are listed in Table S2. Agrobacterium tumefaciens (Agl-0, GV3101 or ABI50) based plant transformation was carried out using the floral dip method (Clough & Bent, 1998). All plant lines examined were homozygous if not indicated otherwise. Homozygosity was determined by antibiotic resistance and 3 independent lines were analysed in the T3 generation.

### Pharmacological treatments

For IAA treatments (10 µM, Sigma-Aldrich, St. Louis, MO) samples were treated for 6h before sampling. TOR inhibitor AZD8055 (10 µM, MedChemExpress, Monmouth Junction, NJ) was applied 16h prior to inducing LR formation by auxin for an additional 6h. Similarly, seedlings were transferred for 16h to 2-Deoxyglucose (20mM, Sigma-Aldrich, St. Louis, MO) containing media to block glycolysis before seedlings were treated for additional 6h with auxin to induce LR formation. Expression of UB10pro>>amiR-TOR was induced via transferring seedlings to plates containing ß-Estradiol (10 µM, Sigma-Aldrich, St. Louis, MO in DMSO) for 24hrs.

### Synchronous induction of lateral root induction

We used the previously described Lateral-root-inducible-system (Himanen *et al*, 2002). In brief, dense horizontal lanes of sterilised seeds were placed on sterile nylon-membranes (SEFAR, Switzerland) and, 7 days after germination, were transferred to plates with fresh ½ MS medium containing 10 µM NPA (Naphthylphthalamic acid, (Sigma-Aldrich, St. Louis, MO) for 24h before shift to 10 µM IAA.

### GC-MS-based metabolite profiling

Profiling of central carbon metabolism intermediates was performed using GC-MS according to metabolite extraction and analysis steps initially described by (Roessner *et al*, 2001). Briefly, 15–40 mg of the previously collected and frozen root tissues were homogenised by tissue lyzer in liquid nitrogen and subsequently mixed with 360 µl ice-cold methanol. 20 µg of Ribitol (Sigma-Aldrich, St. Louis, MO) were added as an internal normalising standard. After extraction (15 min, 70°C), 200 µl chloroform and 400 µl water were added and samples were mixed vigorously before centrifugation. 200 µl of the upper methanol-water phase containing polar to semi-polar metabolites were collected and dried in a vacuum concentrator. Derivatization followed thereafter, including methoximation of the concentrated residues followed by silylation. To this end, the residues were first re-suspended in a methoxyamine-hydrochloride/pyridine solution to methoxymize the carbonyl groups. Samples were then heated (90 min, 37°C) and subsequently silylated with *N*-methyl-*N*-trimethylsilyltrifloracetamide (37°C, 30 min). GC-MS analysis was performed on a gas chromatograph system equipped with a quadrupole mass spectrometer (GC-MS-QP2010, Shimadzu, Duisburg, Germany). For this, 1 µl of each sample was injected in split mode with a split ratio of 1:100 and the separation of derivatized metabolites was carried out on a RTX-5MS column (Restek Corporation, Bellefonte, PA) using instrumental settings optimised by (Lisec *et al*, 2006).

### GC-MS data processing

Raw GC-MS data files were first converted into an ANDI-MS universal file format for spectrum deconvolution and compound identification. Baseline correction, peak identification, retention time (RT) alignment and library matching with the reference collection of the Golm Metabolome Database (GMD, http://gmd.mpimp-golm.mpg.de/) were obtained using the TargetSearch R package from bioconductor (Cuadros-Inostroza *et al*, 2009). Kovats retention indices used for library matching were calculated for deconvoluted mass spectra from measurements of an alkane mixture (Sigma-Aldrich, St. Louis, MO). The Shimadzu GCMS solutions software (v2.72) interface was further used for manual curation of annotation of some metabolites versus authentic standards analysed under the above described analytical conditions. CSV output files (shoot and root data-sets) from the data processing were exported with peak areas obtained for quantifier ions selected for deconvoluted spectra consistently detected in all analysed samples. Peak areas (File S1) were scaled on a sample-basis according to the extracted amount of root tissue and relative to the peak area obtained for the ribitol internal standard in order to correct for putative extraction and analytical performance variations across the different measurements. Finally, peak areas for the above-mentioned compounds obtained in solvent / blank samples were subtracted as background signals from biological samples. For hierarchical clustering analysis of normalized relative peak levels, data were *z*-score transformed and clustering was conducted with the Ward’s clustering method.

### S6K Phosphorylation assay

Proteins were extracted from 40 mg root materials in 200 µl 1X MOPS buffer (0.1M MOPS, 50 mM NaCl, 5% SDS, 10% glycerol, 4 mM EDTA (pH 7.5), 0.3% ß-mercaptoethanol) supplemented with 1.5% phosphatase inhibitor cocktail 2 (Sigma-Aldrich, St. Louis, MO). After adding extraction buffer, samples were briefly mixed and heated at 95°C for 7 min. Cellular debris was removed by centrifugation (10 min, 14,000 rpm, RT). Protein extract were supplemented with 5x Laemmli buffer (Bromophenol blue (0.05%), 0.3 M Tris buffer (pH 6.8), 50% glycerol, 0.1 M DTT) and reheated for 5 min to 95°C. 20 µl protein extract were separated on a 12% SDS gel and transferred to Nitrocellulose membrane (Sigma-Aldrich, St. Louis, MO). Anti-S6K1 (phospho T449) polyclonal antibody (No. ab207399, abcam, Cambridge, UK) was used to detect S6K phosphorylation. S6K1/2 antibody (AS12-1855, Agrisera AB, Vännäs, Sweden) was used to detect total S6K1, Ponceau-S counterstain for confirmation of equal loading.

### Histochemical analysis and microscopy

GUS activity was assayed at 37°C overnight following a modified version of the protocol used in (Weigel & Glazebrook, 2002): the initial washing with the staining buffer (without X-Gluc) and vacuum steps were omitted. GUS staining was followed by fixation in a 4% HCl and 20% methanol solution (15 min at 65°C) followed by 7% NaOH in 60% ethanol (15 min, room temperature). Seedlings were subsequently cleared in successive ethanol baths for 10 mins (40%, 20%, 10%), followed by a 20 min incubation in a solution of 25% glycerol and 5% ethanol. Finally seedlings are mounted in 50% glycerol for imaging with DIC microscopy using an Axio Imager. M1 (Carl 478 Zeiss, Oberkochen, Germany) with a 20X objective. For starch staining, seedlings (18 DAG) were collected in the morning, fixed and cleared as described above, then stained for 30 min with 2 ml of Lugol’s Iodine solution according to (Caspar *et al*, 1985) before visualisation via stereomicroscope (SteREO Discovery.V12, Zeiss, Jena). Calcofluor White counter staining was performed with seedlings fixed for 30 min in 4% PFA in 1X PBS (RT), as described (Ursache *et al*, 2018). Root bend sections were cleared with ClearSee (Kurihara *et al*, 2015) for 1 day and imaged on a Leica SP8 confocal microscope with a 40x, NA = 1.3 oil immersion objective. Calcofluor White fluorescence was detected using the 405 nm excitation laser line, and emission range of 425-475 nm.

### RNA-seq analysis

Samples (Col-0 or inducible *UB10pro>>amiR-TOR* line) were prepared for harvesting using the synchronous induction of lateral root procedure. All samples were pre-treated with 10 µM NPA for 24 h then shifted to plates containing 10 µM NPA and 10 µM Estradiol or DMSO control for additional 24 h before being shifted to either 10 µM IAA + 10 µM Estradiol or DMSO for LR induction or maintained on the same plates. Root tissue was harvested after 6 h. All sampling points were performed in triplicate. For each sample, about 200 segments of the lower two-thirds of the seedling roots were pooled. Total RNA of the 24 samples (2 genotypes x 4 treatments x 3 replicates) was extracted with the Universal RNA purification kit (EURx). Illumina NextSeq libraries were prepared from 2 µg of total RNA and sequencing performed on NextSeq 500 flow cells (12 samples per cell). Reads were mapped onto the *Arabidopsis thaliana* genome (TAIR10) and numbers of reads per transcripts computed using STAR (version 2.5.2b). All the subsequent analysis was done with R (www.r-project.org/) using the DESeq2 package (Love *et al*, 2014). Differentially expressed genes were identified using a grouping variable that combines the Treatment (NPA, NPA_Est, IAA, IAA_ESt) and Genotype (col-0, *UB10pro>>amiR-TOR*) variables at log2 fold change >1 and false discovery rate < 0.05. The procedure was inspired by http://master.bioconductor.org/packages/release/workflows/vignettes/rnaseqGene/inst/doc/rnaseqGene.html#time-course-experiments. The 1141 genes induced by IAA, were defined as the differentially expressed genes (NPA vs. IAA) common that were insensitive to the effect of Estradiol in col-0. The RNASeq data have been deposited to GEO (GSE199202) as part of the SuperSeries GSE199211.

### RT-qPCR

Seedlings were pretreated at 7 DAG for 16h either with 10 µM AZD8055 or DMSO control media before being transferred to DMSO, 10 µM IAA, 10 µM AZD8055 or 10 µM AZD8055 + 10 µM IAA. Root tissues of ca. 200 seedlings were then dissected after 6h. All samplings were performed in quintuplicate. Total RNA was extracted with the RNeasy Plant Mini Kit (Qiagen) and 2µg RNA were reverse transcribed with RevertAid First Strand cDNA Synthesis Kit (Thermo Fisher). Quantitative RT-PCR (RT-qPCR) was performed using gene specific primers (see File S2) in a total volume of 20 μl Absolute qPCR SYBR-green Mix (Thermo Fisher) on a qTOWER^3^ (Analytik Jena) apparatus according to the manufacturer’s instructions.

### TRAP-Seq

The TRAP-seq experiment was conducted according to (Thellmann *et al*, 2020). In brief, seedlings ubiquitously expressing a GFP-tagged RPL18 ribosomal protein (*p35S:HF-GFP-RPL18*, N69096, (Mustroph *et al*, 2009)) were pretreated at 7 DAG for 16h either with 10 µM AZD8055 or DMSO control media before being transferred to DMSO, 10 µM IAA, 10 µM AZD8055 or 10 µM AZD8055 + 10 µM IAA. Root tissues were then dissected after 6h. All sampling were performed in triplicate. For each sample, about 1500 segments of the lower two-thirds of the seedling roots were pooled, flash frozen in liquid nitrogen and later homogenised with a polysome-extraction buffer (Thellmann *et al*, 2020). The suspension was centrifuged for 15 min (16 000 g, 4°C). An aliquot of the homogenate was used for Bulk-RNA-extraction by use of TRIzol reagent (Invitrogen). The remaining supernatant was incubated with GFP-Trap® Magnetic Agarose beads (Chromotek, Munich, Germany). Ribosomal bound RNA was obtained by immunoprecipitation with magnetic anti-GFP beads in accordance with the manufacturer’s instructions and subsequently purified by use of TRIzol reagent (Invitrogen). Illumina NextSeq libraries were prepared from 2 µg of total RNA (bulk) or 100ng (Ribosome bound) after depleting the rRNA via Ribo-Zero Plus rRNA Depletion Kit (Illumina) and sequencing performed on NextSeq 500 flow cells (12 samples per cell). Reads were mapped onto the Arabidopsis thaliana genome (TAIR10) and numbers of reads per transcripts computed using STAR (version 2.5.2b). All the subsequent analysis was done with R (www.r-project.org/) using the DESeq2 package (Love *et al*, 2014). For each assay type (Bulk and TRAP), the variance was stabilised by a r-log transform and z-score were derived for all transcripts genome wide. The log10 ratio of TRAP to Bulk signal was then computed for all transcripts. The auxin responsive genes used for k-means clustering based on the ratio of TRAP to Bulk signal were identified in the Bulk set comparing DMSO to IAA treatment (|log2FC|>1, FDR<0.05). The TRAP-Seq and Bulk RNA seq data have been deposited to GEO (GSE199203) as part of the SuperSeries GSE199211.

### Polysome profile analysis

Root samples were frozen and ground in liquid nitrogen. The powder was resuspended in Polysome extraction buffer [100mM HEPES-KOH pH 8.0, 150 mM KCl, 25 mM Mg(OAc)2, 25 mM EGTA pH 8.0, 0.5% NP-40, 250 mM sucrose, 5 mM dithiothreitol (DTT), Complete Protease Inhibitor EDTA-free (Roche)], and final lysate was cleared by high-speed centrifugation for 15 minutes at 4°C. The equivalent of 100 a.u. (A260; measured on Nanodrop) was layered on top of the 15 to 45% (w/v) sucrose density gradients and then centrifuged at 29,000 rpm in a SW60-Ti rotor for 3 hours at 4°C. The polysome profiles were generated by continuous absorbance measurement at 260 nm using a Gradient Fractionation System (Biocomp Instruments), eight fractions collected, and total RNA from individual fractions was extracted with Tri-Reagent (Trizol) and reverse transcribed with High-Capacity cDNA Reverse Transcription Kit (Applied Biosystem). Quantitative RT-PCR (RT-qPCR) was performed using gene specific primers (see File S2) in a total volume of 10 μl SYBR Green Master mix (Roche) on a LightCycler LC480 apparatus (Roche) according to the manufacturer’s instructions.

### Statistical analysis

All the statistical analyses used in this study were performed in R and Microsoft Excel. The methods and p-values are summarised in the figure legends. Plotting of data was performed using R and Microsoft Excel.

## Supporting information

Stitz et al. 2022 - SuppMat - v1

## Supplementary data

Supplementary data are available online.

## Acknowledgements

We thank A. Leibfreid for critically reading the manuscript, M. Burow (DynaMo Center, Dept. of Plant and Environmental Sciences, University of Copenhagen, Denmark) for *TOR-oe* seeds, S. Savaldi-Goldstein (Faculty of Biology, Technion Haifa, Israel), for the *p35S:HF-GFP-RPL18* line, A. Bishopp (University of Nottingham, UK) for the *pARF19-5’UTR::mVenus* line and Lin Xu (Institute of Plant Physiology and Ecology, Shanghai, PRC) for the *gLBD16-GUS* line and Jazmín Reyes for help with microscopy. The authors gratefully acknowledge the data storage service SDS@hd supported by the Ministry of Science, Research and the Arts Baden-Württemberg (MWK), the COS Metabolomics Core Technology Platform for GC/MS instrument access, the Cluster of Excellence Cellular Networks of the University of Heidelberg (CellNetworks) through grant EcTOP6 “Metabolism and Development” and the German Research Foundation (DFG) through grant INST 35/1314-1 FUGG and INST 35/1503-1 FUGG.

## Author contribution

Conceptualization: MS, EG, AM

Data Acquisition and curation: MS, MR, DK, MSch BB

Formal Analysis: MS, MR, DK, MSch, BB

Funding Acquisition: EG, AM

Methodology: MS, EG, AM

Project Administration: EG, AM

Resources: DJ, AP, JL, AA, RH, MB

Supervision: EG, AM

Visualisation: MS, MSch, EG, AM

Writing – Original Draft: MS, AM

Writing – Review and Editing: MS, DK, MSch, DJ, EG, AM

## Data availability

The data supporting the findings of this study are available from the corresponding author upon request. Sequencing data are available from GEO (SuperSeries GSE199211).

