## Supplementary material for "TOR acts as metabolic gatekeeper for auxin-dependent lateral root initiation in *Arabidopsis thaliana*": Stitz et al. 2022 - SuppMat - v1

### TOR integrates metabolic and developmental cues into Lateral root formation

- File S1. Root and shoot compound relative intensities obtained from the GC-MS metabolomics analysis
- File S2. List of primers used in this study.

- Fig. S1) Glucose and Sucrose levels in shoots of IAA-treated Col-0 and *slr* seedlings.
- Fig. S2) Reconfiguration of the carbon metabolism-related transcriptome during LR formation is influenced by auxin/*slr*-dependent signalling.
- Fig. S3) TOR over activation leads to longer PR roots.
- Fig. S4) Silencing efficiency in *UB10pro>>amiR-TOR* line.
- Fig. S5) Lateral root formation can not be rescued by IAA or external carbohydrate sources in TOR-deficient seedlings.
- Fig. S6) Silencing TOR expression in xylem-pole pericycle cells leads to excessive starch accumulation alongside vasculature.
- Fig. S7) Heatmap showing expression of 475 SLR depend genes 6 hours after using the Lateral Root inducing system in Col-0 and in *UB10pro>>amiR-TOR* root tissues
- Fig. S8) Inhibition of TOR via AZD8055 induces TOR-transcription, but does not reduce transcriptional regulation of key-genes related to lateral root formation.
- Fig. S9) IAA responsive genes detected during ribosome profiling and TOR inhibition IAA induced in the RNA-seq experiment under TOR-deficiency vastly overlap.

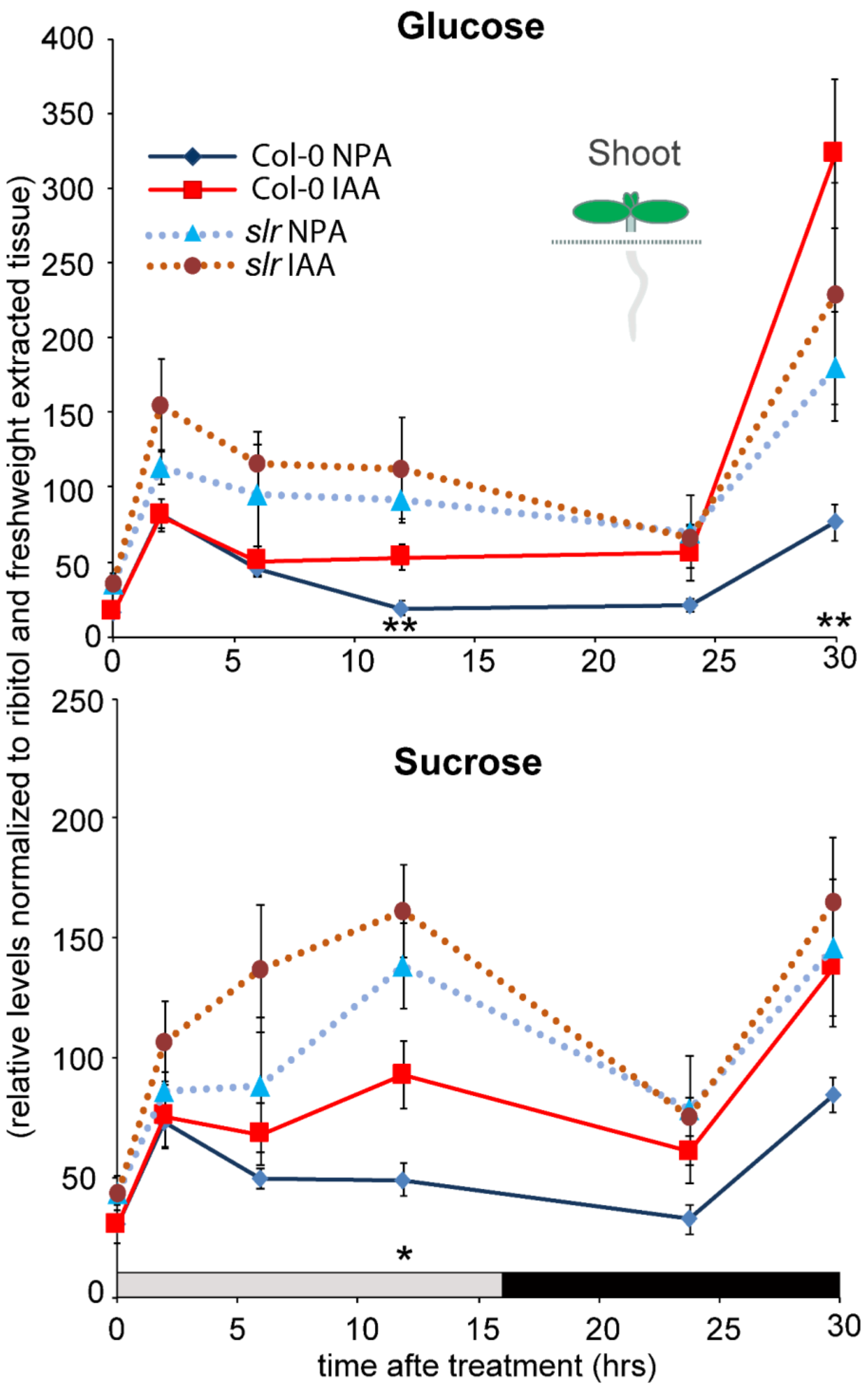


**Fig S1).** Glucose and Sucrose levels in shoots of IAA-treated Col-0 and *slr* seedlings. Mean relative levels (± SE, *n*=5), normalised to the ribitol internal standard and per mg fresh weight) of glucose and sucrose in shoot tissues of Col-0 (solid lines) and *slr* (dashed lines) at the indicated time after IAA application. Asterisks (Col-0), and plus signs (*slr*) indicate significant differences between NPA treated control conditions and auxin (IAA) induced root tissues (unpaired t test; */+ $p < 0.05$, **/++ $p < 0.001$).
Shoot metabolomics data are summarised in File S1.

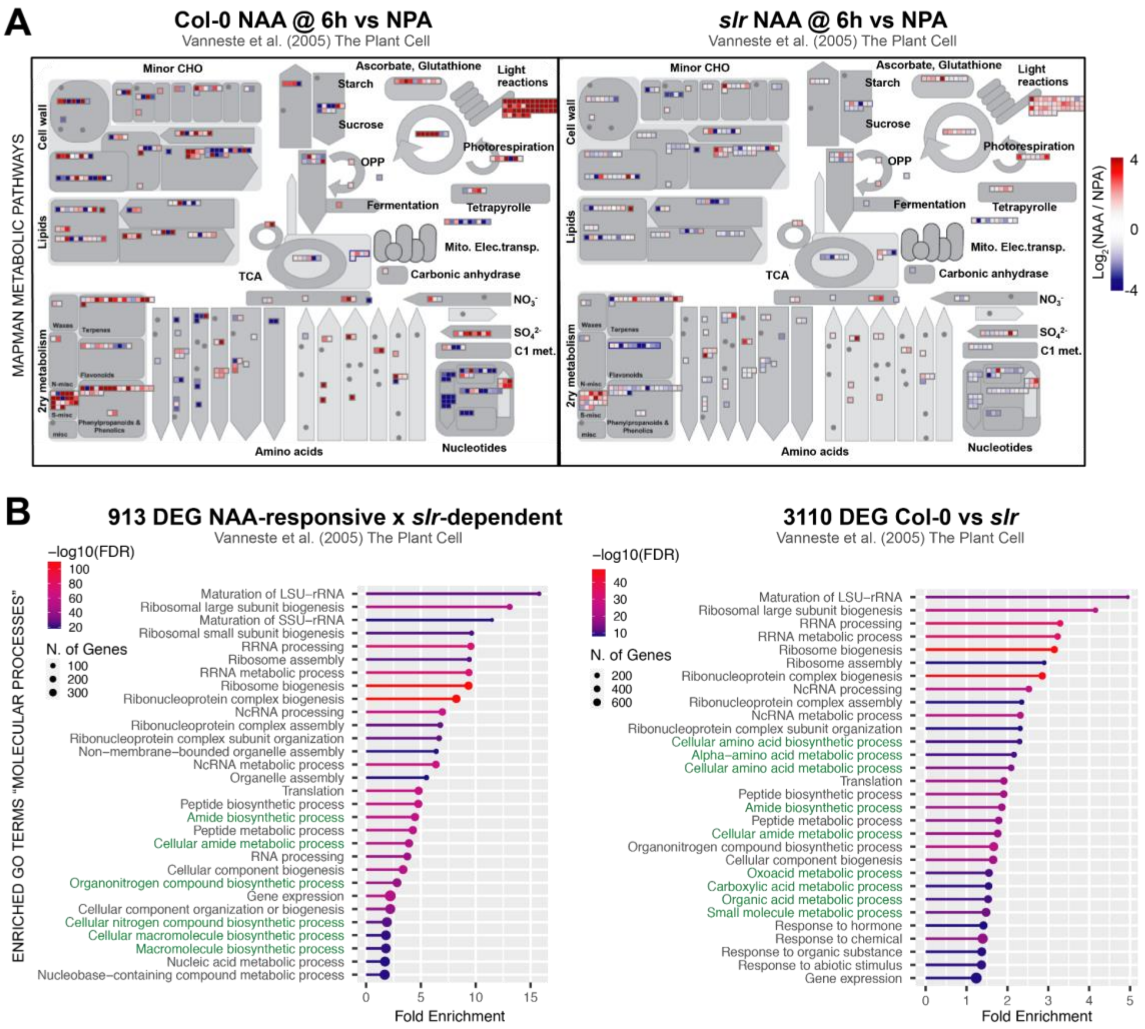


**Fig S2). Reconfiguration of the carbon metabolism-related transcriptome during LR formation is influenced by auxin/*slr*-dependent signalling.**

**A)** MAPMAN-based overview of fold-change ($log_2$-transformed) reconfigurations of central carbon metabolism-related transcripts from root segments of Col-0 and *slr* mutant seedlings after 6h of transfer from NPA to the auxin analog NAA (α-naphthaleneacetic acid). **B)** DEG sets for the comparison of overall Col-0 vs *slr* root transcriptomes (right panel) and specifically associated to an interactive NAA x *slr* effect (left panel) were extracted from the VisuaLRTC transcriptome/statistical data compendium by Parizot *et al.* (2010). Enrichment analyses for Molecular Processes GO terms were conducted using ShinyGO v0.75 (Ge *et al*, 2019) with a FDR P-value cutoff of 0.5 and maximum number of top pathways to show of 30. Central carbon metabolism-related GO terms are highlighted in green. Original transcriptomics data for both analyses are from the Vanneste *et al.* (2005) pioneer exploration of auxin-/*slr*-dependent root transcriptomics responses during synchronized LR induction.

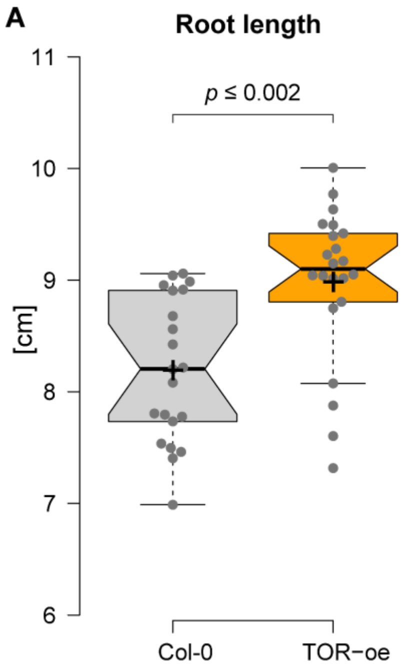


**Fig. S3) TOR over activation leads to longer PR roots.**

14 day old TOR-oe seedlings (GK548) show significantly longer primary roots (A) than Col-0 ($p \leq 0.002$), when grown on ½ MS media containing 2% Sucrose. (unpaired *t*-test).

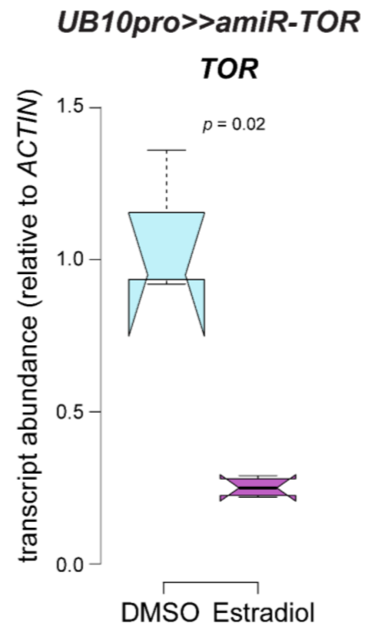


**Fig. S4)** ***TOR*** **silencing in** ***UB10pro>>amiR-TOR***

Root tissues of *UB10pro>>amiR-TOR* plants grown for 24 h on ½ MS media containing 10 µM ß-Estradiol have significantly lower TOR-mRNA levels than *UB10pro>>amiR-TOR* control plants grown for 24 h on ½ MS media containing DMSO control solution (*n*=4). (unpaired *t*-test).

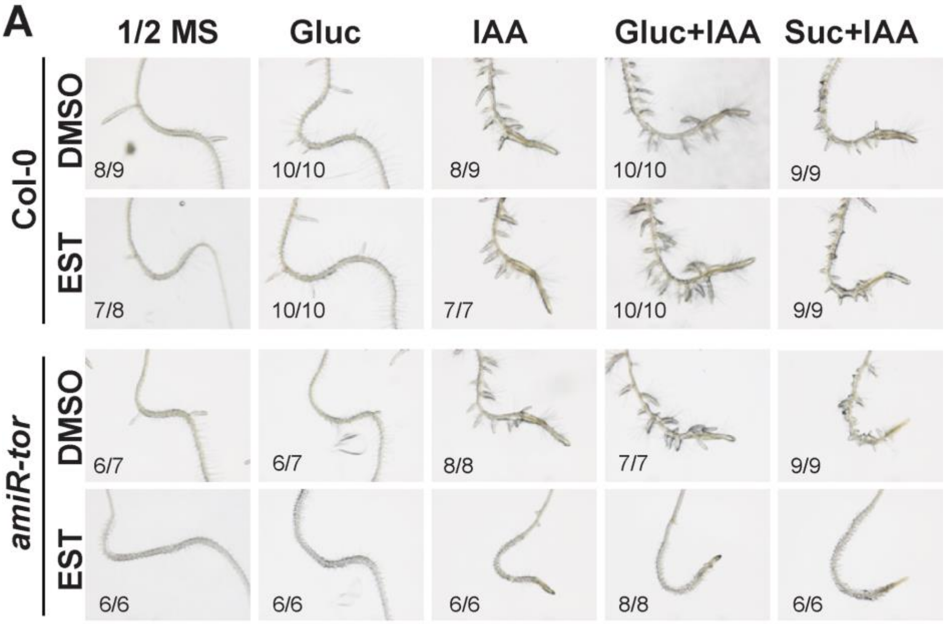


**B**

**Percentage of Rootbends producing LR**

| | Col-0 | | *amiR-tor* | |
|---|---|---|---|---|
| **Treatment** | **DMSO** | **EST** | **DMSO** | **EST** |
| **1/2 MS** | 86 | 88 | 86 | 0 |
| **Gluc** | 100 | 100 | 86 | 0 |
| **IAA** | 100 | 100 | 100 | 0 |
| **Gluc+IAA** | 100 | 100 | 100 | 0 |
| **Suc+IAA** | 100 | 100 | 100 | 0 |

**Fig. S5) Lateral root formation can not be rescued by IAA or external carbohydrate sources in TOR-deficient seedlings.**

**A)** Representative images of Col-0 and *UB10pro>>amiR-TOR* roots after 72h after rescue with 2% Glucose, 10 µM IAA, 2% Glucose + 10 µM IAA or 2% Sucrose + 10 µM IAA. Before transfer to the rescue-media, seedlings were pre-treated for 24h with either DMSO or 10 µM ß-Estradiol. **B)** Quantification of the LR-rescue in UB10pro>>amiR-TOR seedlings.

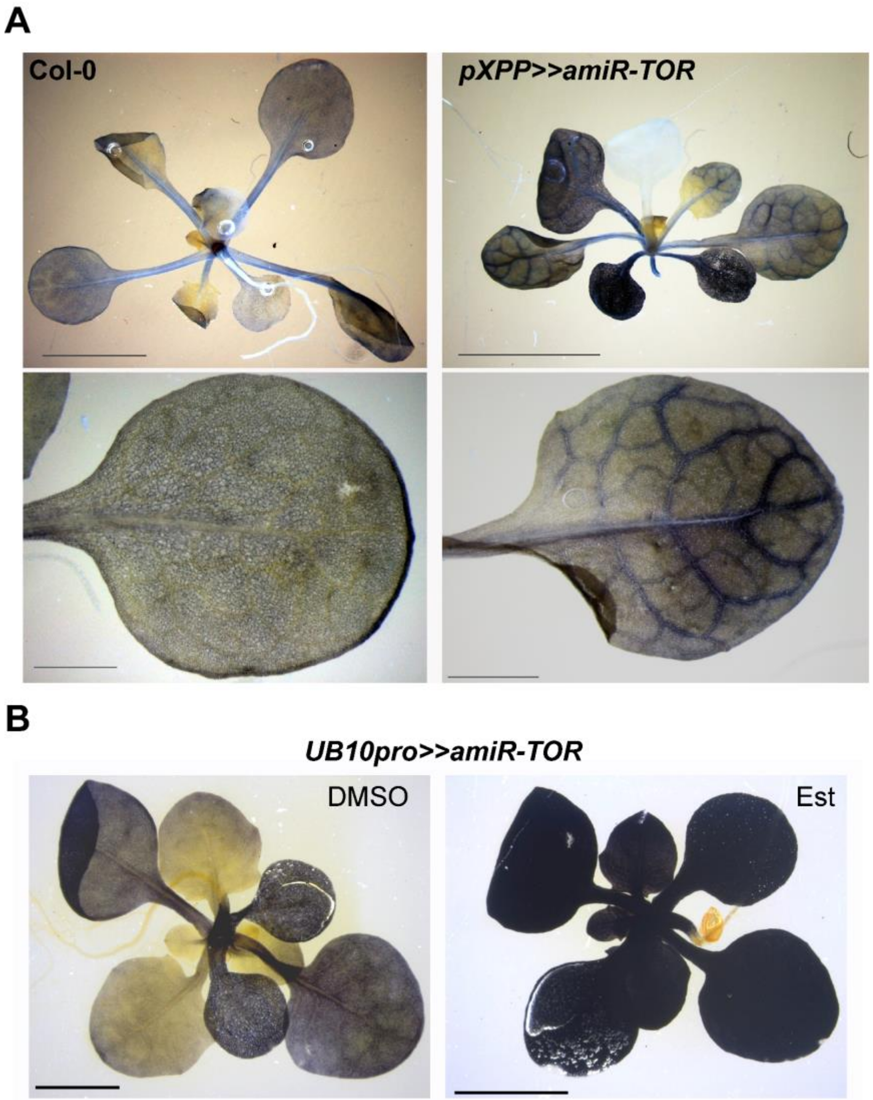


**Fig. S6) Silencing TOR expression in xylem-pole pericycle cells leads to excessive starch accumulation alongside vasculature.**

**A)** In contrast to Col-0, 12 DAG seedlings of *pXPP>>amiR-TOR* plants accumulate starch alongside vasculature of leaves, indicated by pronounced purple coloration around vasculature of DEX grown *pXPP>>amiR-TOR*. (Scale bars: upper panel: 5mm, lower panel: 1 mm). **B)** 14DAG *UB10pro>>amiR-TOR* plants grown for 48hrs on Estradiol accumulate excessive starch throughout the foliage, while DMSO grown *UB10pro>>amiR-TOR* show limited starch accumulations-

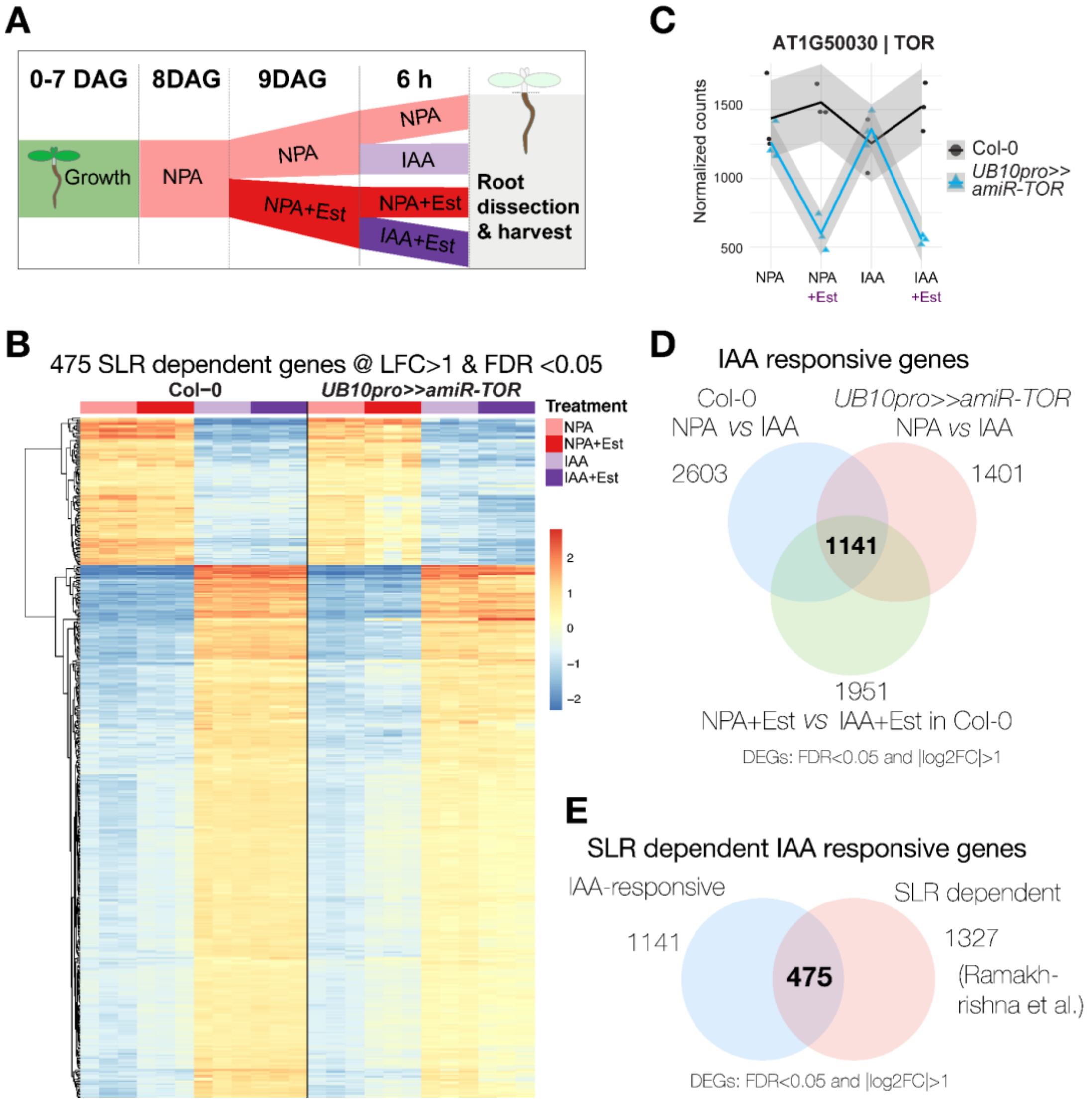


**Fig. S7) Transcriptome analysis upon auxin induced induction of lateral root formation in *UB10pro>>amiR-TOR.***

**A)** Schematic depicting how samples for tis RNA-seq data set were prepared. **B)** Heatmap from a hierarchical clustering analysis (HCA) showing z-score normalised relative levels of 475 SLR dependent genes in tissues +/- induced for LR-formation, and +/- induced for TOR-knockdown (One-way ANOVA, LFC>1 & FDR <0.05) **C)** Extracted traces for RNAseq sample set shows reduction of TOR transcripts in samples generated from UB10pro>>amiR-TOR roots grown on Est containing media compared to control and Col-0 samples. **D)** Venn-diagram of IAA-responsive genes with log 2-fold change >1 f commonly and differentially expressed in Col-0 and *UB10pro>>amiR-TOR* after shifting from NPA to 10 µM IAA, and Col-0 after shift from NPA and Est to IAA and Est. **E)** Venn-diagram showing IAA-responsive genes commonly and differentially expressed in Col-0 and *UB10pro>>amiR-TOR* 6h and Col-0 after shift from NPA and Est to IAA and Est.

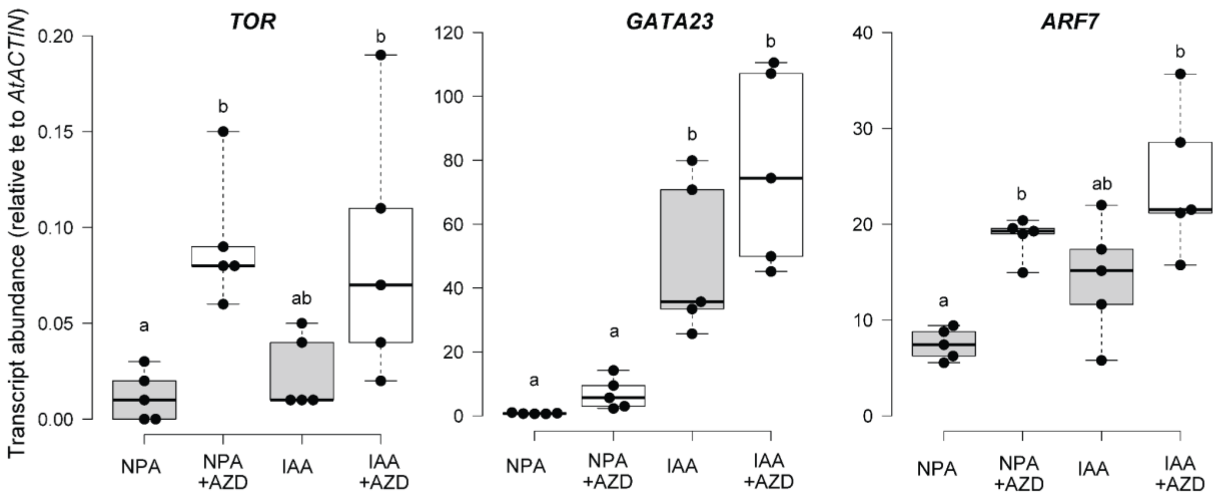


**Fig. S8) Inhibition of TOR via AZD8055 induces TOR-transcription, but does not reduce transcriptional regulation of auxin-induced related to lateral root formation.** Relative expression levels (normalised to *ACTIN*) of TOR are induced by AZD8055, while GATA23 and ARF7 are not reduced by TOR-inhibition. Comparison between samples was performed by one-way ANOVA. Different letters indicate significant differences based on a post-hoc Tukey HSD Test (α = 0.05).

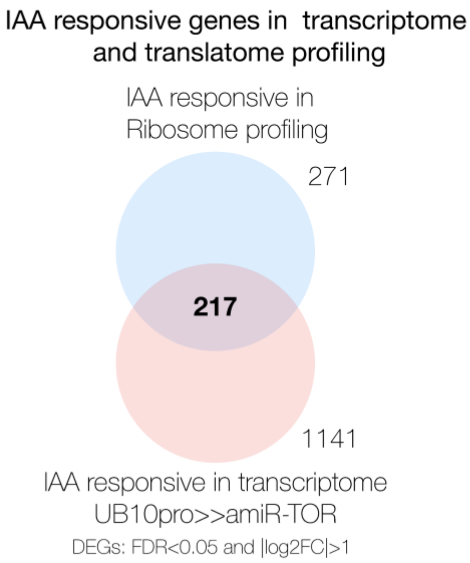


**Fig. S9) IAA responsive genes detected during ribosome profiling and TOR inhibition IAA induced in the RNA-seq experiment under TOR-deficiency vastly overlap.**
Venn-diagram showing IAA-responsive genes commonly and differentially expressed 6h of IAA application in Col-0 after previous AZD8055 inhibition of TOR and, and *UB10pro>>amiR-TOR* on Est.
